# Chemical exhaustion of RPA in cancer treatment

**DOI:** 10.1101/2020.11.30.404640

**Authors:** Pamela S. VanderVere-Carozza, Katherine S. Pawelczak, Navnath S. Gavande, Shadia I. Jalal, Karen E. Pollok, Elmira Ekinci, Joshua Heyza, Steve M. Patrick, John J. Turchi

**Affiliations:** Department of Medicine, Indiana University School of Medicine (IUSM), Indianapolis, Indiana 46202 USA; NERx Biosciences, 212 W 10th St. Suite A480, Indianapolis, Indiana 46202, USA; Department of Pharmaceutical Sciences, Wayne State University College of Pharmacy and Health Sciences, Detroit, Michigan 48201, USA; Herman B Wells Center for Pediatric Research, Departments of Pediatrics, Pharmacology & Toxicology, Medical & Molecular Genetics Indiana University Simon Comprehensive Cancer Center, Indianapolis, Indiana 46202, USA; Department of Oncology, Wayne State University School of Medicine and Barbara Ann Karmanos Cancer Institute, Detroit, MI 48201; Department of Biochemistry and Molecular Biology, Indiana University School of Medicine, Indianapolis, Indiana 46202, USA

**Keywords:** Replication protein A, DNA damage response, cancer therapeutics, lung cancer, chemical exhaustion, combination therapy

## Abstract

Replication protein A (RPA) plays essential roles in DNA replication, repair, recombination and the DNA-damage response (DDR). We have developed second generation RPA inhibitors (RPAi’s) that block the RPA-DNA interaction. These DNA-binding inhibitors (DBi’s) can elicit a state of cellular RPA exhaustion resulting in single agent *in vitro* anticancer activity across a broad spectrum of cancers and *in vivo* activity in two non-small cell lung cancer models. The cellular response to RPAi treatment suggests a threshold exists before RPA inhibition induces cell death. Chemical RPA exhaustion potentiates the anticancer activity of other DDR inhibitors as well as traditional DNA damaging cancer therapeutics. Consistent with the chemical RPA exhaustion model, we demonstrate that the effects of RPAi on replication fork dynamics and DNA damage signaling are similar to other known DDR inhibitors. In accordance with the RPA threshold model, retrospective analysis of lung cancer patient data demonstrates high RPA expression as a negative prognostic biomarker for overall survival in smoking-related lung cancers. Similarly, relative expression of RPA is a predictive marker for response to chemotherapy. These observations are consistent with the increase in RPA expression serving as an adaptive mechanism that allows tolerance of the genotoxic stress resulting from carcinogen exposure. These data demonstrate a unique mechanism of action of RPAi’s eliciting a state of RPA exhaustion that impacts the DDR and may provide an effective therapeutic option for difficult to treat lung cancers.

**Graphical Abstract:** 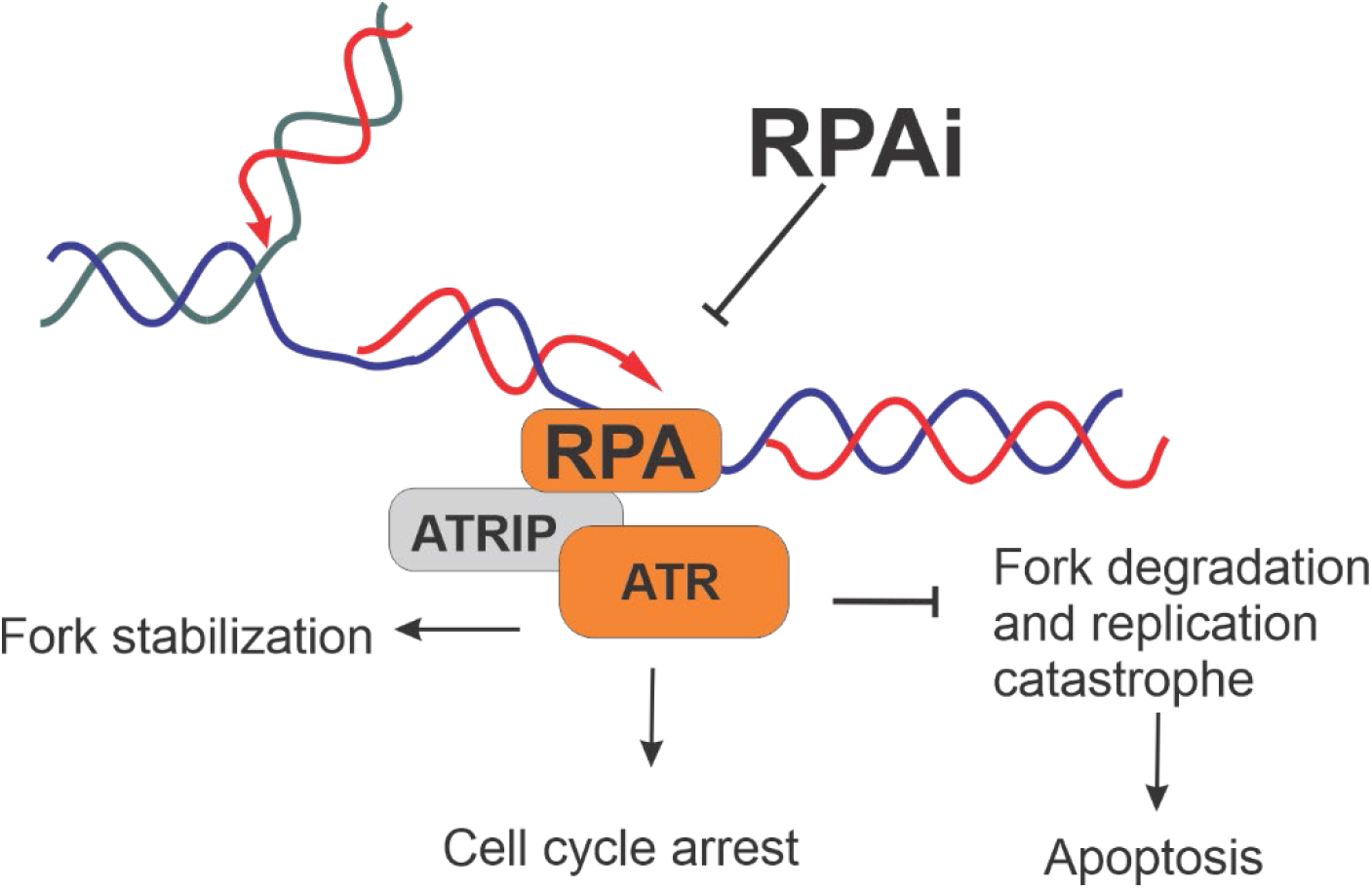

## 1. Introduction

Recent advances in kinase targeted agents and immuno-oncology (IO) therapy have changed many treatment paradigms for lung cancer, however, the majority of the over 200,000 individuals diagnosed each year in the US alone will die from their disease [1]. Late diagnosis, early metastasis and co-morbidities all contribute to the poor prognosis of most lung cancer patients. Lung epithelial cells are exposed to a variety of carcinogens that can be dramatically increased in cigarette smoke exposure, and likely contributes to the high mutation burden observed in smoking related cancers. It stands to reason that lung epithelium would have a robust DNA repair capacity to counter the DNA damage elicited by cigarette smoking and early research demonstrated the importance of DNA repair in lung carcinogenesis [2;3]. This repair capacity can explain the rapid resistance to cancer therapeutic modalities that induce DNA damage including two frequently used platinum-based agents, cisplatin and carboplatin, and ionizing radiation [4]. Recent advances in our understanding of how cells, both normal and cancerous, respond to DNA damage stress has identified a number of unique vulnerabilities that can be exploited for effective therapy to treat cancer [5;6].

The DNA damage response (DDR) is the coordinated activation of several pathways that sense DNA damage or replication stress. Important sensors of DNA damage include the three upstream DDR kinases: (i) DNA-dependent protein kinase (DNA-PK), (ii) Ataxia telangiectasia mutated (ATM) and (iii) ATM-Rad3 related kinase (ATR). Downstream signaling from these DDR kinases includes activation of effectors that regulate DNA metabolic processes, cell cycle progression and cell survival [7–9]. Each kinase employs a unique DNA binding subunit to initiate DDR activation, DNA-PK senses DNA breaks via the Ku 70/80 dimer binding to ends of DNA, whereas ATM senses DNA DSBs via its DNA binding subunit MRN (Mre11/Rad50/Nbs1). In contrast, ATR is activated by single-stranded DNA via RPA and ATRIP rather than DNA breaks [10]. Single-stranded DNA is an intermediate in many normal cellular process and can be exacerbated during DNA replication stress, excessive DNA damage and dysregulated transcription [11;12]. RPA binding protects single-stranded DNA from degradation and serves as the platform for ATRIP and ATR binding. However, kinase activation remains low until a kinase conformational change is induced by TOPBP1 or ETAA1 [13;14]. ATR activation in response to DNA damage and replication stress serves to activate the CHK1 kinase in response to DNA replication stress and subsequently, cdc25 phosphatase to impact cell cycle progression and also to suppress replication origin firing to limit or delay replication initiation events. All of these events are coordinated to maintain genomic stability and the fidelity of chromosomal duplication. Mechanistically, inhibition of ATR in combination with DNA damage has been demonstrated to induce replication fork collapse, chromosomal pulverization and cell death. Based on these data, several ATR inhibitors have been developed and are now in early phase clinical trials as monotherapy or as part of multimodal regimens [15;16].

RPA plays a central role in the initiation of the ATR signaling pathway which offers a unique mechanism for regulating the DDR. Adequate levels of RPA are necessary to ensure any single-stranded DNA generated is adequately protected from degradation [17;18]. The partial depletion of RPA via siRNA increases sensitivity to DNA damage and induces replication stress and catastrophe, similar to the effect of ATR inhibition. This “RPA exhaustion” renders cells vulnerable to the replication stress or DNA damage associated with cancer growth and progression. We hypothesize the inhibition of the RPA-DNA interaction can induce a state of chemical RPA exhaustion and lead to an increased susceptibility to DNA damage, thus enhancing the therapeutic window for existing chemotherapy and targeted therapies. In this report, we demonstrate that first-in-class small molecule chemical inhibitors of RPA can reduce the active pool of RPA and impact the DDR. These effects translate to both single agent anti-cancer activity and synergy with a number of DNA-targeted therapeutics. These data coupled with retrospective clinical data identify a subset of high RPA expressing lung cancer patients that could benefit from RPA targeted therapy.

## 2. Results

### 2.1 Retrospective analysis of RPA expression in lung cancer

Considering the model of RPA exhaustion and the potential that the expression of RPA could impact the DDR, we sought to determine how the expression of RPA impacted survival in lung cancer. We selected lung cancer as lung epithelial cells are continuously exposed to a wide array of potentially carcinogenic agents, a situation exacerbated by smoking and second-hand smoke exposure. To assess the potential clinical utility of RPA inhibition we performed a retrospective analysis of gene expression data in lung cancer as a function of smoking history and response to chemotherapy treatment. In current and former smokers, the data reveal that high RPA expression is a negative prognostic biomarker correlating with worse overall survival (Figure 1a). This difference in survival as a function of RPA expression was also observed when selecting for patients that received adjuvant chemotherapy (Figure 1b), suggesting that low RPA is predictive of a better response. In the analysis of never smokers (Figure 1c) no correlation between RPA expression and survival was observed. Importantly, this population is likely a collection of heterogeneous cancer phenotypes which is characterized by a higher level of driver mutations in growth signaling pathways and as such, these never smokers are expected to have received targeted kinase inhibitor therapy. The finding that RPA expression level does not impact survival is therefore not surprising. Collectively these data suggest that potential genotoxic damage induced by smoke exposure induces reliance on RPA expression to protect against genotoxic stress that, if reversed, could impact survival.

**Figure 1.**
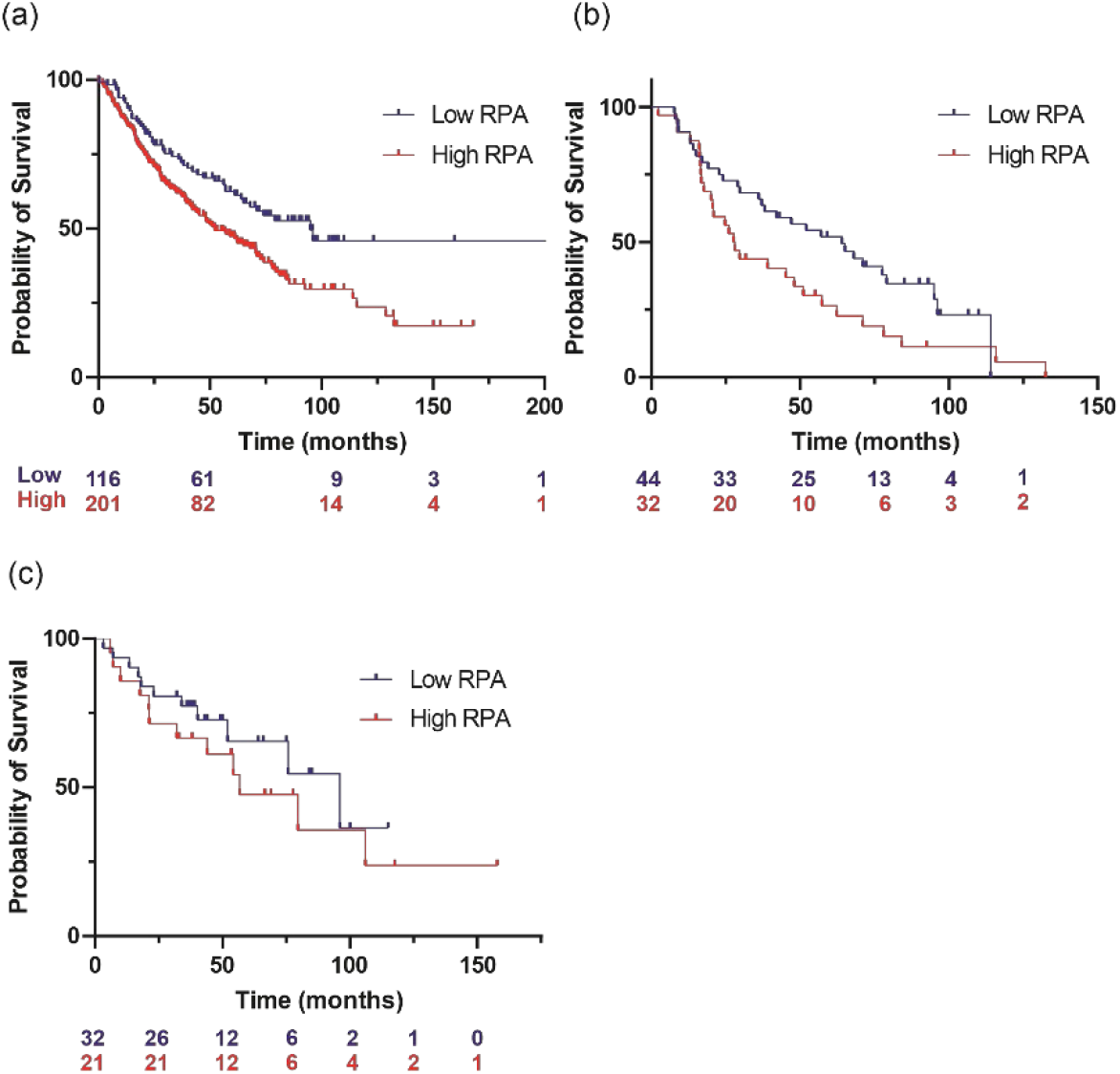
Retrospective analysis of overall survival as a function of RPA expression in NSCLC. (a) Former and current smokers. (b) Former and current smokers that received chemotherapy. (c) Never smokers.

### 2.1. Chemical RPA exhaustion

Our previous analyses of reversible RPAi’s revealed chemical liabilities that limited their utility in cell-based assays and *in vivo* [19;20]. We have further optimized the **NERx-551** candidate to generate candidate RPAi’s **NERx-329** and -**2004** (Figure 2a) that possess potent RPA inhibitory activity (Figure 2b). The data also show the compounds are specific for inhibiting the RPA ssDNA interaction as the interaction of E. Coli SSB with ssDNA (indicated by the asterisk) is not impacted by the RPAi’s. These compounds also display excellent solubility, cellular uptake and physicochemical properties [21]. As the addition of a propylmorpholino to the oxopentoic acid moiety increased solubility and cellular uptake we sought to assess single agent cellular anticancer activity in the H460 NSCLC cell line (Figure 2c). The data demonstrate that both compounds **329** and **2004** possess potent single agent activity compared to their **551** predecessor as assessed by CCK-8 metabolic assay. The RPAi mechanism of action proposed would be expected to be tumor-agnostic, and we therefore predicted that RPAi would display activity in a variety of tumor types. We therefore assessed RPAi activity in other cancer models including ovarian cancer (A2780), testicular cancer (GCT27) and a NSCLC adenocarcinoma (A549) (Figure 1d). The effective cell killing was observed across these diverse cancer cell types. These cell lines and cancer models were selected as they all represent cancers that initially respond to DNA damaging chemotherapy. Interestingly, the titration curves obtained with the different cell lines displayed different response across cell lines, with both the A2780 and GCT27 cell lines having steep slopes with Hill-coefficients less than −10 while A549 and H460 were less steep with a corresponding Hill-coefficient value of ~5. These data suggest the presence of a threshold whereby low doses are essentially nontoxic, but after reaching the threshold exposure, cells are inviable. These findings are consistent with the model of RPA protection of ssDNA as a mechanism to maintain genome stability.

**Figure 2.**
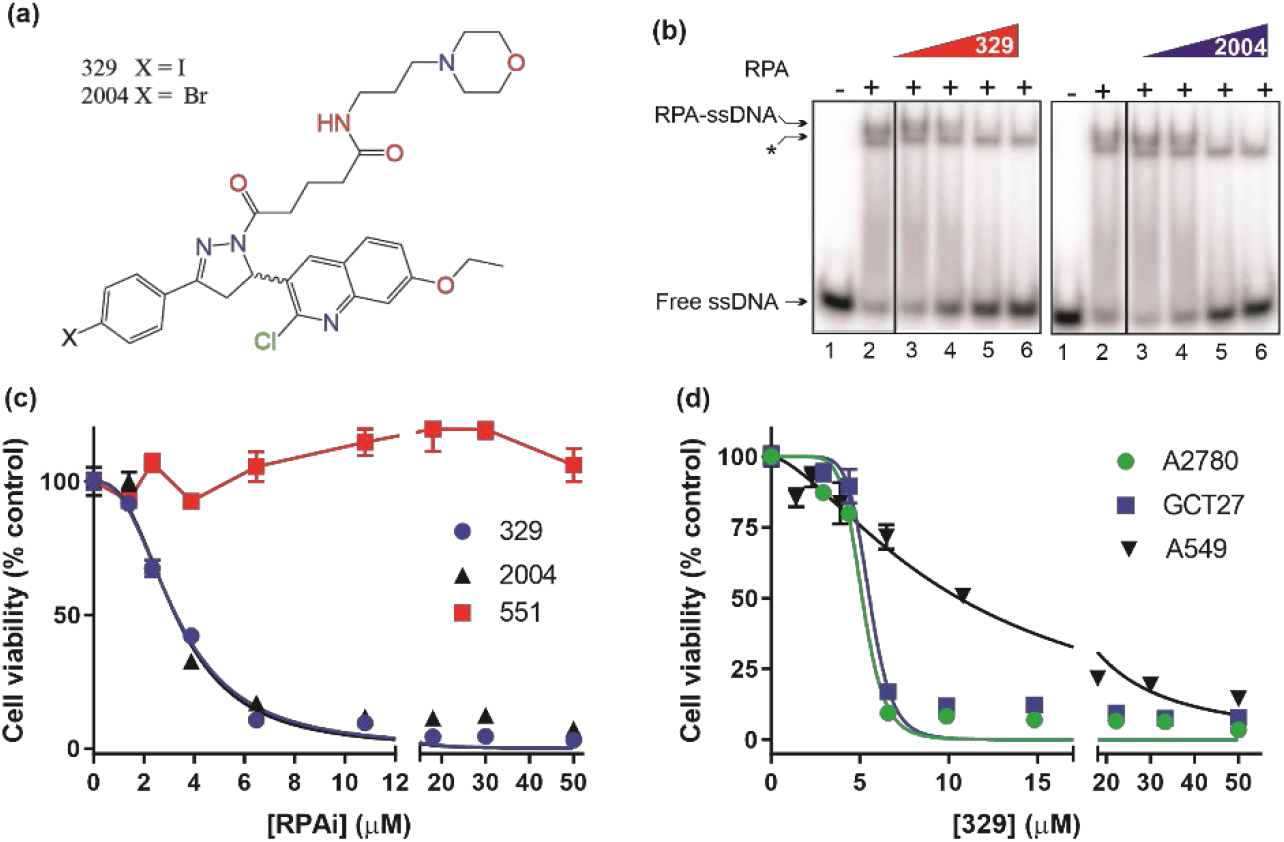
RPAi inhibitory activity. (a) Chemical Structure of RPAi’s **329** and **2004**. (b) EMSA analysis of RPA-DNA interaction inhibition by 329 and 2004. Lane 3-6 in each panel contain 6.25 12.5.25 and 50 μM of the indicated RPAi respectively. The * indicates the position of the E. coli SSB-ssDNA complex that serves as an internal specificity control (c) Cell viability of H460 NSCLC cells in response to **329** and **2004**. (d) Cell viability of A2780, GCT27 and A549 cancer cell lines as a function of **329**.

Analyses of single agent activity of compound **329** in 60 discrete cancer cell lines across a variety of solid tumors revealed similar findings. A range of IC_50_ values were obtained with most falling between 5 and 10 μM and largely independent of tumor site (Figure 3a). Certain uterine, lung and esophageal cancer cells lines were the most sensitive while pancreatic adenocarcinomas tended to be more resistant. Interestingly, the Hill coefficients on the other hand spanned a much larger range (Figure 3B), which did not necessarily correlate with the potency as measured by IC_50_. Certain lung, muscle, ovarian, and cervical cancer lines were characterized by the lowest Hill coefficients. These data are consistent with the tumor agnostic nature of RPA inhibition and a mechanism of action involving a threshold of measurable cytotoxic sensitivity. Two cell lines, A2780 and A549 were also included in the 60-cell line screen that were presented in Figure 2d. In each assay the A2780 cells display a reduced hill coefficient compared to the A549. The absolute difference between the two is less in the 60-cell screen., This is likely the result of the different treatment times and viability assays used with the 60-cell line screen using a 72-hour incubation with RPAI and the measure of ATP as a readout of cell number/viability while the data in Figure 2 were performed with a 48-hour incubation of RPA 1 and the CCK-8 assay for viability which assess mitochondrial dehydrogenase activity.

**Figure 3.**
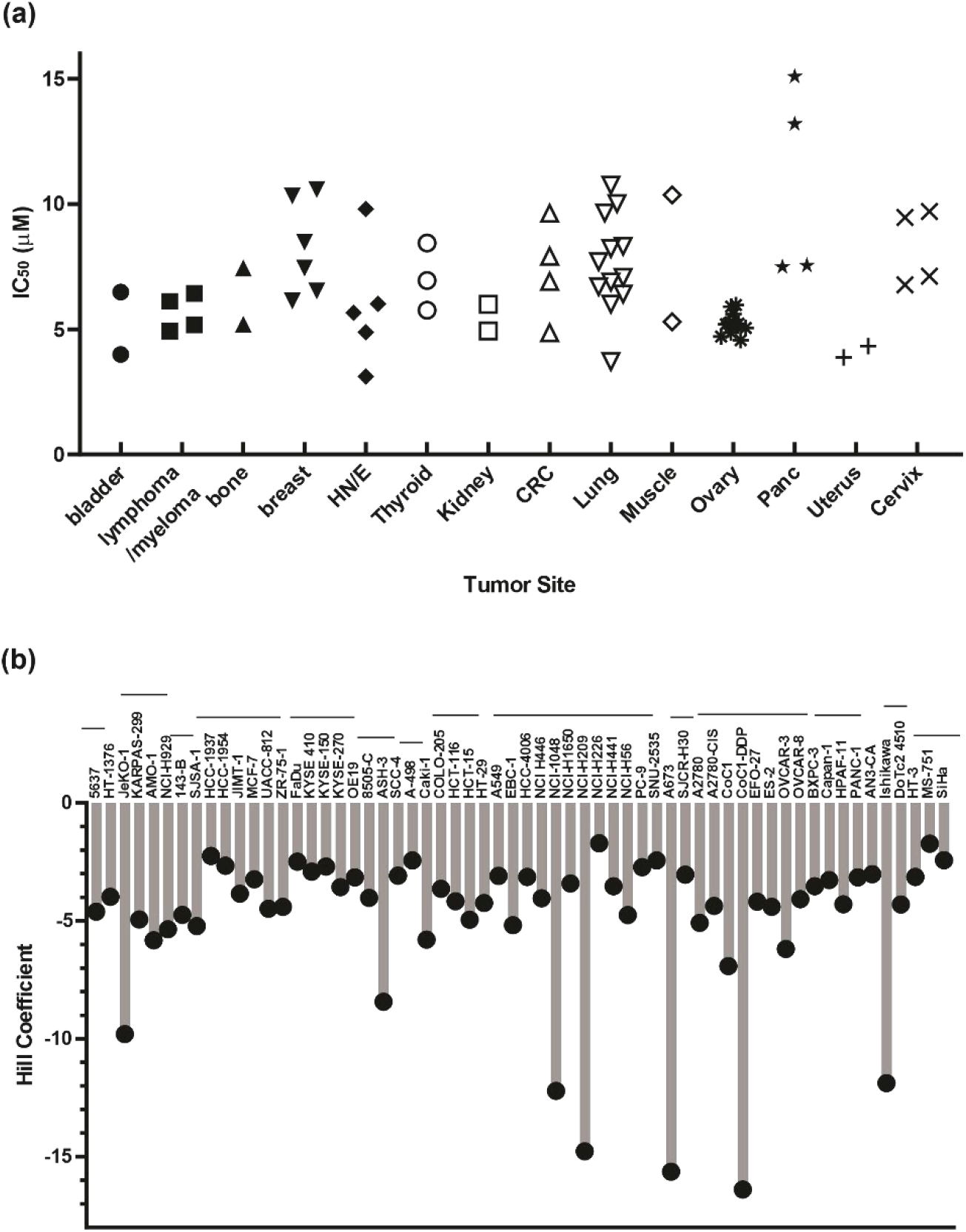
Cellular activity of 329 in 60 cancer cell lines. Cell lines were treated with a 4-log range of RPAi 329 for 72 hours. Cell viability was assessed using CellTiter-Glo luminescent viability assay. The data represent the average of triplicate treatments and the data were fit using non-linear regression analysis to calculate cellular IC_50_s. (a) IC_50_ results from each cell line grouped by tumor type. (b) Hill coefficients for individual cell lines. The horizontal lines above cell line names indicated the tumor sites in the order depicted in panel (a). Complete data for each cell line is in Supplemental Table 1.

Predecessor reversible RPA inhibitors **505** and **551** also displayed single agent anticancer activity, although this was not accompanied by caspase activation or annexin V/PI positivity suggesting a non-apoptotic mechanism of cell death [19]. The increased cellular uptake and potency displayed by the morpholino containing compound **329** prompted us to revisit this activity. Using the activation of caspase 3 and 7 as a readout, we demonstrate that **329** induces cell death via a classical apoptotic pathway (Figure 4) and the activation of caspase 3 and 7 clearly distinguishes it from predecessor compound **551**. The titration analyses assessing apoptosis correlated with the corresponding CCK-8 viability curves, and show the presence of a modest threshold. The apoptosis assays were conducted after a 24 hours drug treatment while CCK-8 viability assays require 48 hours for maximal effect. Assessment of apoptosis at 48 hour was similar to 24 hours in terms of the titration though the maximum signal detected was higher, as expected.

**Figure 4.**
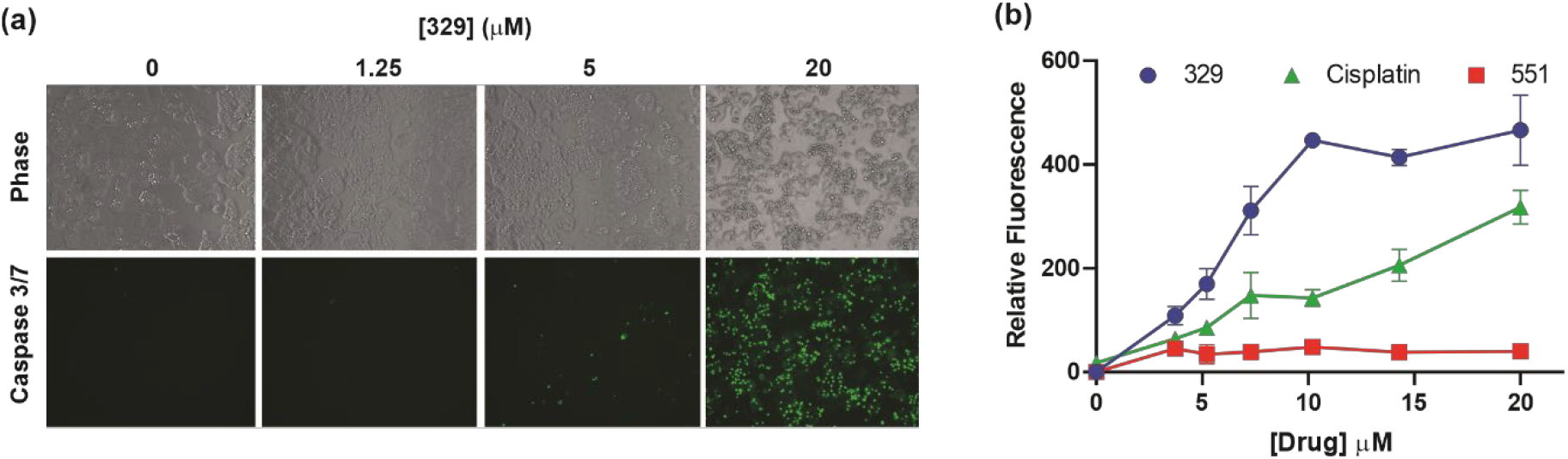
329 induction of apoptotic cell death. (a) Analysis of caspase3/7 activity in H460 cells following 24 hours of treatment with 1 % DMSO or the indicated concentrations of **329**. Fluorescence images were captured as described in Methods. (b) Quantification of caspase 3/7 activity. Fluorescence was measured in 96 well plates using a Biotec Synergy 1 plate reader following 24-hour incubation with the indicated drugs and concentrations.

### 2.2 In vivo activity

The long-term goal is to move towards efficacious and safe combination therapies that include RPA inhibition. To this end, we conducted single-agent screening in two lung cancer cell line-derived xenograft models. Predecessors to **329** and **2004**, compounds **505** and **551** possessed modest *in vivo* activity [20]. Having optimized cellular uptake and solubility *via* the addition of the propyl morpholino, we sought to determine how these modifications impact *in vivo* anticancer activity using two NSCLC models, A549 adenocarcinoma and H460 large cell carcinoma. Analysis of toxicity revealed that safe dosing could be achieved up to 200 mg/kg with no overt toxicity and no significant loss in body weight similar to predecessor compounds [20]. Assessment of liver and kidney function also showed no differences from vehicle controls. We observed potent single agent anticancer activity in both models with differing dosing regimens.

Considering the rapid growth kinetics of H460, we administered **2004** and **329** at 50 mg/kg twice daily for three days (Figure 5a). With this dosing strategy, a decrease in tumor volume was observed with **2004** at day 10 indicative a not just a slowing of tumor growth by tumor shrinkage. Compound **329,** on the other hand, showed slower tumor growth. Continued daily dosing of **329** at 50 mg/kg did not result in a statistically significant reduction in tumor weight, a result that was obtained with continued **2004** (Figure 5a and 5b). *In vivo* efficacy data obtained in the slower growing A549 model revealed that daily dosing of **329** at 200 mg/kg resulted in a significant reduction in tumor volume (Figure 5c) while 2004 at 50 mg/kg led to a trend of decreasing tumor volume that was not statistically significant. Together these data demonstrate the utility of RPAi in treating lung cancer and further analysis including combinations will require optimized formulation, dosing and schedule to achieve maximal anti-cancer activity.

**Figure 5.**
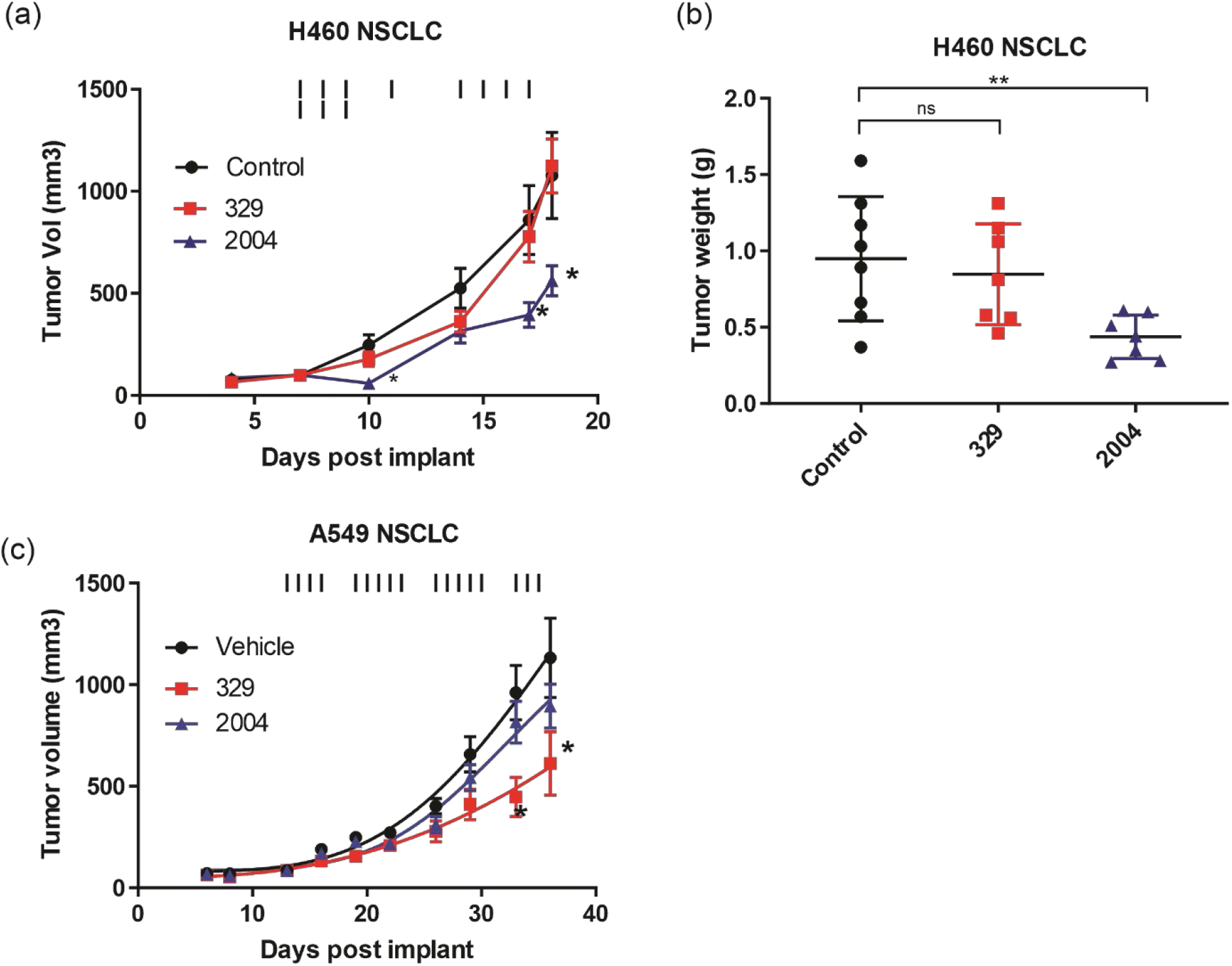
In vivo analysis of RPAi 329 and 2004. (a) H460 NSCLC cells were implanted subcutaneously, mice were randomized and treatment initiated at day 7. Vehicle, RPAi **2004** or **329** (50 mg/kg) was administered IP as indicated by the vertical lines. Tumor volume was measured by calipers and (b) tumor weight determined at day 19. (c) A549 cells were implanted subcutaneously, mice were randomized and treatment initiated at day 10. Compounds **329** (200 mg/kg) and 2004 (50 mg/kg) was delivered IP for 3 weeks, with a schedule of 5 days on and 2 days off. Tumor volume was measured by calipers. Statistically significance difference from vehicle treated tumors are indicated by the asterisk *=p<0.05; **p<0.01

### 2.3 Therapeutic combinations

Considering RPA’s role in numerous DNA metabolic processes, we determine how inhibition of RPA impacts sensitivity to a variety of DNA damaging chemotherapeutics that induce different types of damage. Interestingly, we observe synergy with agents that cause replication stress, bulky lesions, and DNA double strand breaks (Figure 6a), whereas no synergy was observed with paclitaxel, a non-DNA damaging therapeutic. These results suggest that the cytotoxic effects of RPAi’s may be mediated by a broader effect on the DNA damage response as opposed to suppression of individual replication and repair pathways. This is supported by data demonstrating that the rates of removal of cisplatin from DNA is not appreciably altered in RPAi treated cells (Figure 6b). This effect was observed in p53 wildtype H460 NSCLC cells as well as the p53 null H1299 NSCLC cell line, though the difference in initial damage was greater in the H460 cells. Considering this data, we suspected that RPA inhibition would synergize with other DDR targeted therapeutics to block multiple pathways within the more broadly concerted DDR. We therefore assessed synergy of RPAi’s with a series of DDR targeted agents that are currently in clinical trials (Figure 6c). The data demonstrate modest synergy is observed with each agent in the H460 NSCLC cell model with exception of the Wee1 inhibitor. Interestingly, we did observe modest synergy with the PARPi BMN637 in BRCA wild type cells despite the relatively limited activity seen with single agent PARPi in these cells. Not surprisingly, we have demonstrated a greater degree of synergy between RPAi and PARPi in BRCA1 null cells compared to BRCA wild type cells [22]. Interestingly, both ATR and DNA-PK inhibition were more effective when used in combination with RPAi treatment suggesting that either inhibition of parallel pathways as in the case of DNA-PK or sequential inhibition of a single pathway in the case of ATR, contributes to enhanced increased anticancer activity. Wee1 inhibition however was antagonistic or additive with RPAi over the entire range of cells affected which places its activity down stream of RPA as expected.

**Figure 6.**
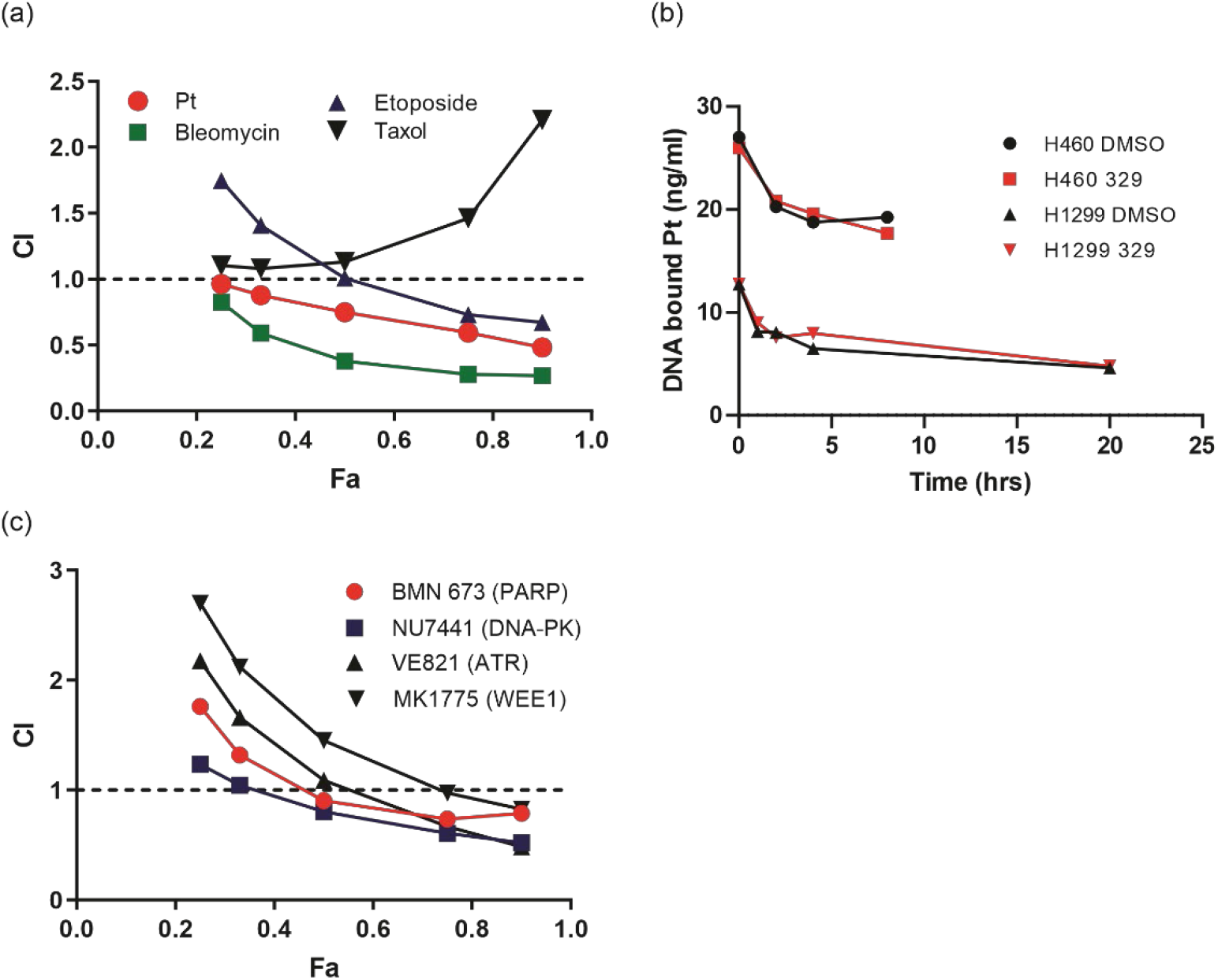
Analysis of RPAi 329 combination treatment. (a) Chou-Talalay analysis of combination with chemotherapeutics. The combination index (CI) is plotted as a function of the fraction of cell affected (Fa) for each treatment combination the **329** (b) Repair of cisplatin lesions as a function **329** as measured by ICP-MS. (c) Chou-Talalay analysis of combination with DDR targeted agents as described in panel (a).

### 2.4 Mechanism of RPAi-Induced Suppression of the DDR

The data demonstrating both single agent activity and synergy with agents that cause differing type of DNA damage suggest that RPA inhibition might be influencing the global DNA damage response. As RPA is required for sensing ss-DNA and signaling via ATRIP to ATR, we posit that chemical inhibition of RPA would block the DDR. To assess this possibility, we have assessed the DDR by measuring replication fork dynamics and markers of DDR activation as a function of RPA inhibition. We employed quantitative image-based cytometry to assess the DDR measuring γ-H2AX and chromatin bound RPA. Experiments were conducted in H460 cells treated with hydroxyurea (HU) to induce replication stress. As a control, cells were concurrently treated with HU and VE821, an ATR inhibitor. This combination has been shown to induce RPA exhaustion and replication catastrophe [17;18]. We then assessed the effect of RPA inhibition in similar experiments. The data presented in Figure 7 demonstrate minimal effects on γ-H2AX and RPA loading as a function of single agent RPA inhibition or ATR inhibition. This is consistent with a reduction but not ablation of RPA binding capacity. Replication stress induced by HU-dependent fork stalling results in increases in γ-H2AX and RPA loading as expected. The combination of HU and ATR inhibition showed increased γ-H2AX as expected and also a significant increase in RPA loading consistent with the hyper-RPA loading from DDR inhibition. The combination of HU and **329** gave the increased γ-H2AX levels consistent with DDR blockade, and an increase in RPA loading that was less than that observed with ATRi. These data are consistent with the chemical exhaustion of RPA by **329** but could also be explained by the decreased potency of 329 versus ATRi. The similar responses to replication stress between ATRi and RPAi are also suggestive of actin in the same pathway as would be expected considering RPA’s crucial role ATR activation[23] Replotting the data a function of cell cycle, determined by DAPI intensity, allows observation of the alterations in S-phase cells (Figure 7c). This presentation highlights the difference in H2Ax levels between RPAi and ATRi while the number of cells with hyper loaded RPA is les in RPAi treated cells than in ATRi treated cells.

**Figure 7.**
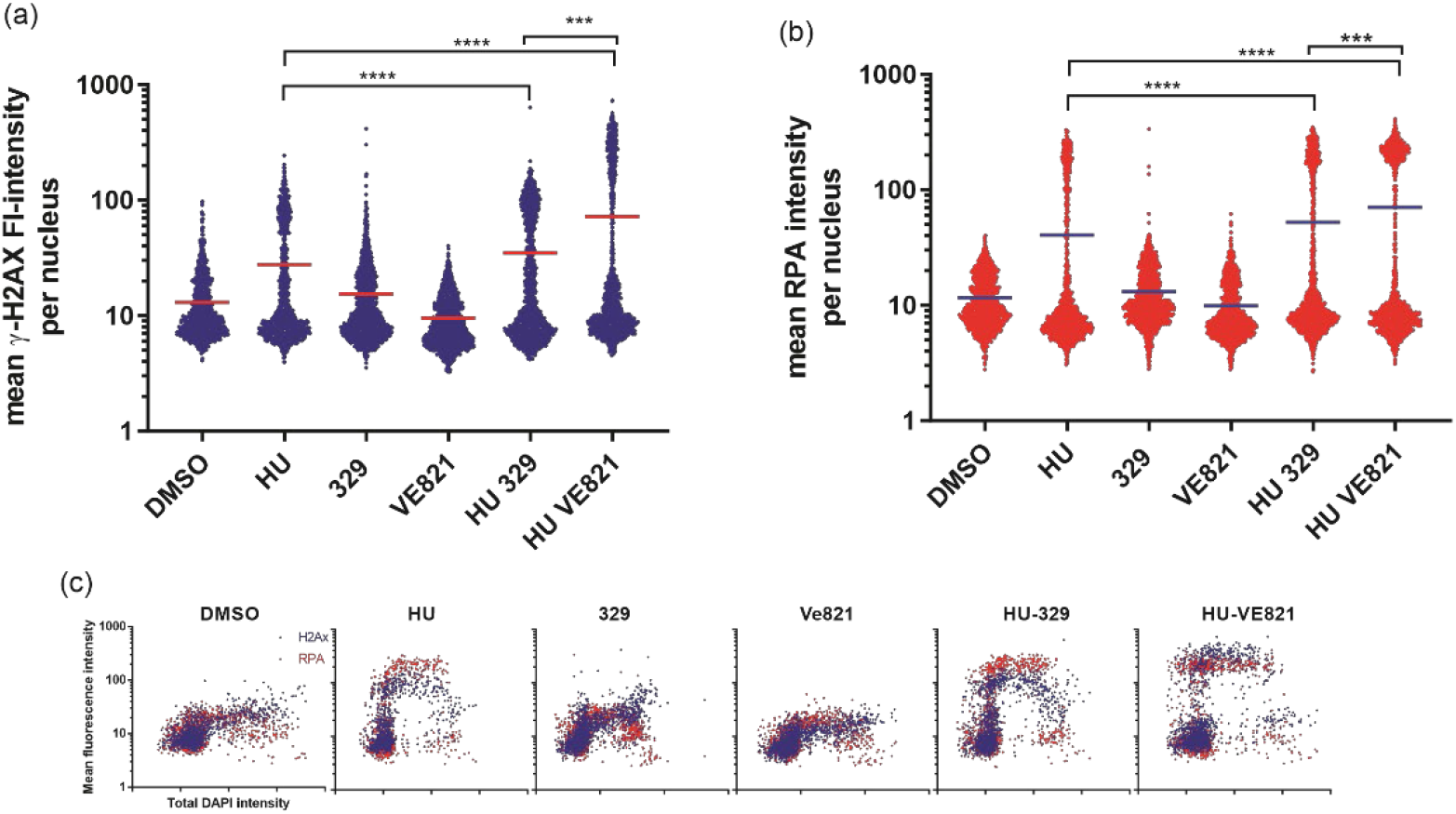
RPAi impact on DDR. H460 NSCLC cells were treated with the indicated agents and DDR activation measured by γH2AX (a) and RPA foci formation (b). Mean fluorescence intensity was determined from captured images by ImageJfrom 200 individual cells per treatment. Statistical significance is indicated **** p<0.0001, *** p<0.001. (c) Replot of the data as a function of total DAPI intensity to indicate cell cycle position.

A further measure of altered DDR induced by RPA inhibition is the degradation of replication forks upon stalling and RPA exhaustion. We therefore assessed replication fork dynamics and nascent strand degradation using DNA fiber analysis. The treatment scheme is depicted in Figure 8a. We first pulse-labeled replicating DNA with IdU for 20 minutes. After IdU removal, replication forks were stalled by the addition of HU or left to replicate with vehicle treatment. HU was removed and replication labeled with CldU. Following CldU cells were treated with the DDRi or vehicle. The data obtained are presented in Figure 8b. Representative images are presented in Supplemental Table 2. As expected minimal effects were observed with ATRi or RPAi alone. However, in cells that received HU and then either ATRi or RPAi, a significant decrease in CldU/IdU signal was observed. This data suggests that the addition of DDRi after fork stalling by HU results in nascent strand degradation at stalled replication forks. Importantly, the effect of RPAi was similar to ATRi as expected for targets in the same pathway. Representative images are presented in Figure 8c. These data suggest that DDR checkpoint abrogation by ATRi or RPAi and a subsequent increase in the presence of unprotected ssDNA in S-phase results in replication fork instability and nascent strand degradation.

**Figure 8.**
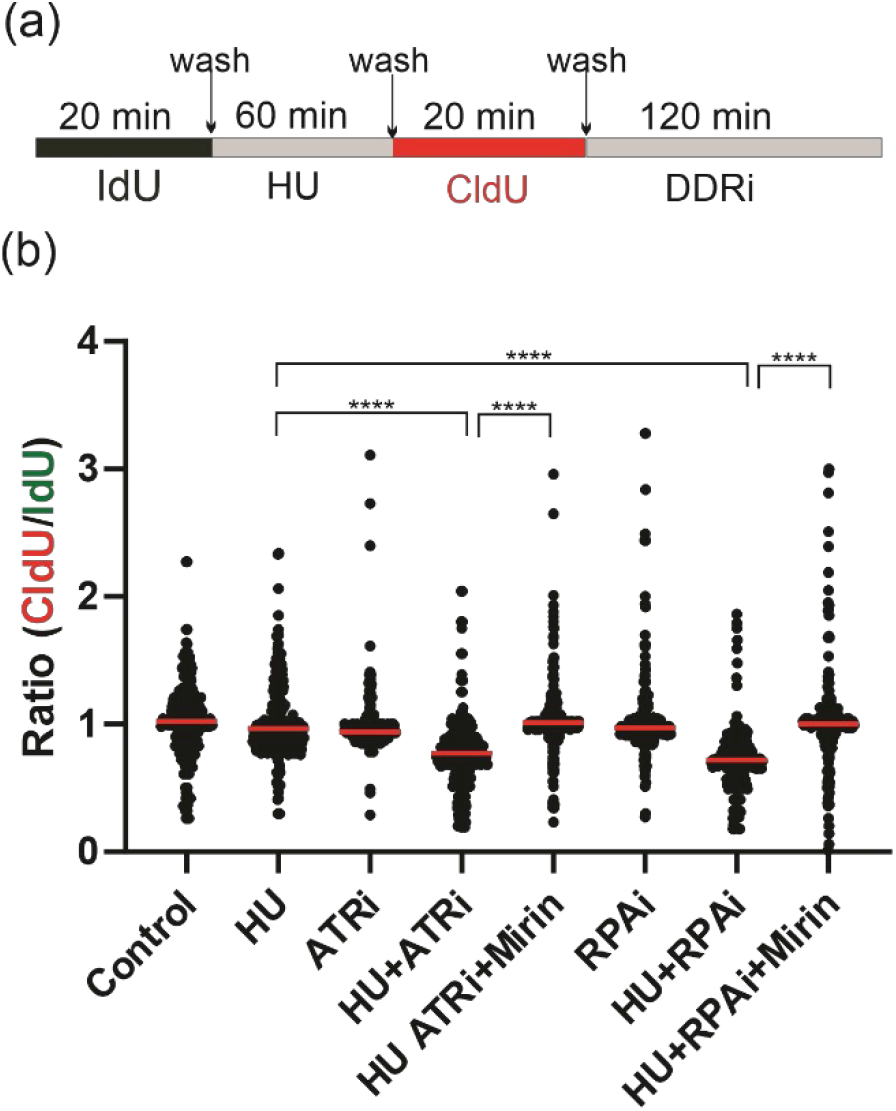
RPAi impact replication fork dynamics. (a) Schematic depiction of experimental design. (b) Quantification of results from DNA fiber analysis in H460 cells treated with the indicated agents. HU was used at a final concentration of 2.5 mM, the ATRi VE-822 at 2 μM and the RPAi NERx-**329** at 50 μM. Data presented are combined from three individual experiments (100 fibers analyzed per experiment; 300fibers total). Red bar indicates the median value of CldU/IdU. Data were analyzed by ANOVA with Bonferroni test for multiple comparisons (****p<0.0001).

## 3. Discussion

The DNA damage response is actively being pursued for cancer therapy with phase I results being recently reported for ATR inhibitors [15]. The vast majority of individual targets being developed in the DDR space are kinases, largely a function of the advances made over the past decade on developing kinase targeted agents in the growth signaling pathways [24]. Kinases, however represent a minority of the protein components in the DDR pathway and larger replication, repair and recombination pathways [8;25]. There are myriad opportunities to impede the DDR via non-kinase targeted agents [26–28]. The DDR pathway is initiated by sensing DNA discontinuities, damage or DNA structures via DNA binding modules associated with each kinase DNA-PK, ATM and ATR. We have a targeted these unique protein-DNA interactions with small molecules to first elucidate specific mechanisms of DDR activation that can be used to guide the development of cancer therapeutics [21;26;29–31]. RPA is a complex target as a function of its roles in multiple DNA metabolic and catabolic pathways [32]. Two classes of RPAi’s were initially discovered: (i) covalent RPA modification agents and (ii) reversible inhibitors that target the oligonucleotide/oligosaccharide binding folds (OB-folds) responsible for the RPA-DNA interaction [33]. In this report using optimized reversible RPAi’s we demonstrate single agent *in vivo* activity and synergy in combination with traditional and DDR targeted therapy. Furthermore, we probed the putative mechanism of RPAi’s anticancer activity.

PARPi therapy has now been approved in 4 different solid tumors with prostate and pancreatic joining ovarian and breast in the list of approved indications. Recent evidence on the mechanism of PARPi suggests that single stranded DNA and specifically lagging strand gaps, contribute to PARP efficacy [34]. If this mechanism is relevant one could envision that BRCA wild type cells would be sensitized to PARPi if the DDR was chemically inhibited. Our finding of synergy as measured by Chou-Talalay combination index analysis support this basic finding and extends to our recent analyses in BRCA1 deficient cells which show BRCA1 deficient cells are hypersensitive to RPAi compared to BRCA complemented cells [22]. The observation of synergy in BRCA wild type cells indicates suggests that RPA inhibition could impair homologous recombination repair (HRR) to create an HR phenotype that increases the potency of PARPi. The impact on HR could be in addition to the effect on the DDR. This result is consistent with our observation of synergy with both ATR and DNA-PK inhibition which can be explained by RPAi impacts on individual, parallel pathways or cross talk between the DDR signaling events. An alternative hypothesis is that another aspect of RPA involvement could explain the synergy, including an alteration in replication fork stability and restart. This is supported by the single molecule studies that the effects of RPAi on replication dynamics are similar to that of ATRi effects which remains consistent with the dependent nature of ATR on RPA DNA binding activity in signaling replication fork stress. It is interesting that ATR activity is impacted by ATM as well based on recent studies in both *in vitro* models and patient responses in clinical trial data. This suggests that cross talk between the three arms of the DDR, DNA-PK, ATM and ATR, is advantageous if not necessary for responding to replication stress or DNA damage. The ability to block the binding of RPA to single stranded DNA can induce differential effects depending on the RPA requirement for each pathway. For instance, the amount of RPA needed for nucleotide excision repair (NER) of cisplatin treated cells is anticipated to be very low based on the cellular levels of cisplatin damage. Accordingly, our observation that RPAi does not impact NER catalyzed repair is not surprising. Similarly, in normal, unperturbed DNA replication RPAi has minimal effects on our initial assessment of replication dynamics, however, when fork stalling is induced by HU, a dramatic effect of RPAi is observed, consistent with the increase in the amount of RPA needed to address the replication stress and the limited RPA available as a function of the inhibitor.

The model of RPA threshold is consistent with our analysis of RPAi cellular activity and the tumor agnostic mechanism of action. Also consistent with these data are previous findings that RPA expression has been described as a prognostic and predictive biomarker in a small number of studies [35–38]. Our retrospective analysis of NSCLC confirms and extends these studies to demonstrate that RPA expression levels can be both prognostic and predictive in smoking associated lung cancers. Its role in the DDR is likely critical to respond and protect from the myriad of genetic insults stemming from carcinogen exposure. It is therefore interesting to speculate that RPA expression or activity may also be predictive of response to other DDR targeted therapeutics.

## 4. Materials and Methods

**RPA inhibitors-** RPAi **329** and **2004** were synthesized and characterized as previously described [21].

EMSA: EMSAs were performed as previously described [21]. Briefly, reactions were conducted in 20 mM HEPES (pH 7.8), 1 mM DTT, 0.001% NP-40, 50 mM NaCl. RPAi’s were suspended in 100% DMSO and DMSO concentration in the final reaction mixture was constant at less than 5%. Purified full length RPA (120 ng) was incubated with the indicated RPAi or vehicle in reaction buffer for 30 min before the addition of the [^32^]P-labeled 34-base ssDNA probe. Reactions were incubated for 5 min at room temperature and products separated via 6% native polyacrylamide gel electrophoresis. The bound and unbound fractions were quantified by phosphor-imager analysis using ImageQuant software (Molecular Dynamics, CA) and data fit by non-linear regression using GraphPad Prism.

### CCK-8 viability assays

Cell lines were obtained from ATCC and maintained as monolayer cultures in RPMI medium (H460, GCT27, A2780) or DMEM (A549) supplemented with 10% fetal bovine serum. H460 and A549 cells were plated at 2.5 x 10^3^ and A2780 and GCT27 cells plated at 5 × 10^3^ were plated in wells of a 96-well plate and incubated for 18-24 hours prior to treatments. Cells were treated with the indicated concentration of RPAi for 48 hours. The vehicle (DMSO) concentration was held constant at 1%. Cell metabolism/viability was assessed by a mitochondrial metabolism assay (CCK-8) as we have described previously [26]. The generation of the water-soluble formazan product by cellular dehydrogenases is proportional to the number of living cells. Following incubation with CCK-8 reagent, absorbance was measured at 450 nm with a BioTek Synergy H1 plate reader. Values were compared to vehicle-treated controls to determine percent viability and the results represent the average and SEM of triplicate determinations.

### Apoptosis assay

Apoptosis induction was determined by activation of Caspase 3 and 7 using the CellEvent™ Caspase-3/7 Green Detection Reagent (Invitrogen).H460 cells were plated at 5 × 10^3^ cells/well in black 96 well plates with clear bottoms (Costar) and incubated for 24 hours prior to treatments. Cells were treated with the indicated concentration of RPAi or cisplatin for 24 hours. The vehicle (DMSO) concentration was held constant at 1% for RPAi treatments. For caspase 3/7 detection, media was removed and replaced with PBS containing 5% FBS and 2uM CellEvent™ Caspase-3/7 Green Detection Reagent. Cells were incubated at 37°C/5% CO2 for 1 hour and fluorescence intensity was measured in a BioTek Synergy H1 plate reader (excitation/emission 485/528). Images were captured with an Evos FL2 Auto microscope (Invitrogen) using a 10X objective.

### Cell viability in 60 cancer cell lines

90μL cell suspensions were seeded in 96-well plates in respective culture medium with a final cell density of 4×10^3^ cells/well and incubated overnight. 10× solution of **329** (top working concentration: 40 μM of test article in media with 3.16-fold serial dilutions to achieve 9 dose levels) was prepared and 10μL of drug solution or culture medium containing 0.5% DMSO (vehicle control) was added to plate (triplicate for each drug concentration). Plates were incubated for 72hr at 37°C with 5% CO2 and then measured by CellTiter-Glo assay (Promega). Briefly, plates were equilibrated at room temperature for 30 min, and 50 μL of CellTiter-Glo reagent was added to each well. Contents were mixed for 5 min on an orbital shaker to induce cell lysis, and plates were further incubated at room temperature for 20 min to stabilize the luminescent signal. Luminescence was recorded using EnVision Multi Plate Reader. Percent cell growth was calculated relative to DMSO treated cells (vehicle control) and the data were fit using non-linear regression analysis (PRISM GraphPad) to calculate cellular IC_50_.

### DNA Fiber Analysis

Analysis of DNA replication intermediates was performed as previously described with minor modifications [39;40]. H460 cells were seeded in 6-well plates at a density of 2 x10^5^ cells. The following day, cells were labeled with IdU (20 μM) for 20 minutes, followed by treatment with HU (2.5 mM) for 60 minutes, then released into CldU (200 μM) for 20 minutes, followed by treatment with ATRi (2 μM, Selleckchem, S7102) for 2 hours or RPAi (50 μM, **329**) for 2 hours. After harvesting the cells were resuspended in PBS at a concentration of 1,000,000 cells/mL, 2 μL of the cell suspension was mixed with 8 μL of lysis buffer (200 mM Tris-HCl pH 7.5, 50 mM EDTA, 0.5% SDS) on a Superfrost Plus microscope slide (Fisher Scientific). After 6 minutes of incubation, the slides were tilted at a 45-degree angle to allow cell lysates to slowly run down the slide. After air-drying, the slides were fixed in methanol: acetic acid (3:1) and stored at 4°C. DNA fibers were denatured with 2.5N HCl for 1 hour, washed with PBS, and blocked with 5% BSA in PBS-T (PBS + 0.1% Tween-20) for 1 hour. DNA fibers were incubated with rat anti-BrdU antibody (1:50, Abcam, ab6326) for CldU and mouse anti-BrdU antibody (1:50, BD Biosciences, 347580) for IdU in a humid chamber at 37°C for 1 hour. After washing, slides were incubated with secondary antibodies (anti-rat Alexa 488, 1:100) and anti-mouse Alexa 568 (1:100) at room temperature for 45 min. Excess antibodies were removed by washing with PBS-T 3 times. After air-drying, the slides were mounted onto a coverslip with mounting medium. Fiber tracts were imaged with a Nikon epifluorescence microscope using a 40x oil immersion objective and 100 fibers for each group were analyzed in ImageJ where the ratios of CldU: IdU were compared using pixel length. Data were analyzed by ANOVA with Bonferroni test for multiple comparisons.

### Combination studies

To assess synergy, the combination index (CI) was determined as described by Chou-Talalay as we have previously described [20]. Briefly, H460 cells were treated with RPA inhibitor and the indicated agent alone and in combination. The range of treatment was dependent on the IC_50_ of each agent and the range was ¼ to 3 x IC_50_. The data from both the single agent treatments as well as the combination treatment were used to calculate the CI and plot this value as a function of the fraction of cells affected (Fa). A CI of > 1 indicates antagonism between the two agents, while a CI < 1 indicates synergy. A CI of 1 demonstrates an additive effect.

### Repair of Cisplatin-DNA adducts

Repair of Pt-DNA damage was determined by measuring the DNA-bound Pt via ICP-MS as we have previously described [41]. Cells were treated with cisplatin and the indicate RPAi and at the indicated times after, DNA was isolated, quantified, and hydrolyzed in 1% nitric acid at 70°C overnight. The amount of Pt per ug of DNA was determined by ICP-MS analysis.

### *In vivo* analyses

To assess anti-cancer efficacy the hind flanks of sixty 8-10 week old NSG mice were implanted with the indicated cells (~2 x 10^6^) in matrigel. Tumor volumes were monitored by electronic caliper measurement [tumor volumes = length x (perpendicular width)^2^ x 0.5]. NSG studies were approved by the Institutional Animal Care and Use Committee at Indiana University School of Medicine. Male Nod SCID gamma (NSG) (NOD-scid/IL2Rg^null^) mice (In Vivo Therapeutics Core Facility, IU Simon Comprehensive Cancer Center, Indianapolis, IN, USA) were used and housed in a pathogen-free facility at IUSM LARC. Mice with tumors of approximately 100 mm^3^ were randomized into individual treatment arms. The indicated RPAi was formulated in 10% DMSO Tween 80 and administered via intraperitoneal injection (IP) at the indicated times. Tumor volumes were monitored biweekly as indicated and the results are presented as the average tumor volume ± standard error of the mean for each group. The number (n) for each experiment is presented in the figure legend.

## 5. Conclusions

In conclusion, this study demonstrates the utility of RPA inhibition as a cancer therapeutic strategy. We demonstrate the ability to identify design and optimize small molecule inhibitors of the RPA protein-single stranded DNA interaction and deploy these to impinge on the DDR pathway. Inhibition of the DDR pathway results in single agent anticancer activity and synergy with agents that either induce DNA damage or block other steps in the cellular response to DNA damage. Mechanistic studies reveal that RPAi act similar to ATRi in DR activation and replication fork dynamics, placing them in the same pathway. Interestingly, we do observe synergy between RPAi and ATRi suggesting on-target effects of RPA inhibition could be impacting other DNA metabolic pathways. These data support a model where a threshold level of RPA is required to support and protect DNA stability and maintenance. Chemical inhibition of the RPA-DNA interaction can induce a state of RPA exhaustion rendering the cells unable to cope with the stress and ultimately induce cell death.

## Author Contributions

Conceptualization, J.T. and P.V.C.; methodology, J.T., P.V.C., K.E.P., S.J.; validation, J.T. and S.M.P.; formal analysis, J.T. S.M.P. and S.J.; investigation, P.V.C., K.S.P. N.G. J.H. and E.E.; resources, J.T. K.S.P.; data curation, J.T. K.E.P. S.M.P. K.S.P.; writing–original draft preparation, J.T..; writing–review and editing, P.V.C., K.S.P. N.G. S.M.P. J.H. E.E. K.E.P.; visualization, J.T. S.M.P. K.E.P. P.V.C.; supervision, J.T. S.M.P.; project administration, J.T..; funding acquisition,.J.T. K.S.P. and S.M.P. All authors have read and agreed to the published version of the manuscript.

## Acknowledgments

We thank Tony Sinn and the IUSCCC In Vivo Therapeutics Core Facility which generated the NSG mice used in this study and conducted the *in vivo* studies.

