## Supplemental Tables for "Chemical exhaustion of RPA in cancer treatment"

Table 1. 60 cell line **329** sensitivity parameters

| Cell line | Hill coefficient | EC50 |
|---|---|---|
| NCI H446 | 4.96 | -4.04 |
| SiHa | 7.19 | -2.43 |
| FaDu | 8.09 | -2.49 |
| EBC-1 | 6.21 | -5.19 |
| SJCR-H30 | 5.06 | -3.04 |
| NCI-H56 | 7.05 | -4.76 |
| A2780 | 5.05 | -5.09 |
| SNU-2535 | 5.63 | -2.44 |
| KYSE-270 | 2.93 | -3.57 |
| HT-1376 | 6.17 | -3.99 |
| HPAF-11 | 7.29 | -4.30 |
| "5637 | 4.26 | -4.61 |
| CoC1 | 5.21 | -6.92 |
| ASH-3 | 6.87 | -8.43 |
| A-498 | 4.66 | -2.44 |
| HT-3 | 6.77 | -3.14 |
| AN3-CA | 3.71 | -3.03 |
| OVCAR-8 | 4.39 | -4.08 |
| PC-9 | 6.22 | -2.72 |
| Ishikawa* | 3.99 | -15.28 |
| OVCAR-3 | 5.82 | -6.19 |
| EFO-27 | 5.36 | -4.19 |
| JIMT-1 | 8.79 | -3.85 |
| A549 | 6.76 | -3.10 |
| HCC-1954 | 7.47 | -2.66 |
| A2780-CIS | 4.41 | -4.37 |
| DoTc2 4510 | 7.04 | -4.30 |
| 8505-C | 5.57 | -4.03 |
| HCC-1937 | 7.02 | -2.25 |
| NCI-H226 | 8.76 | -1.71 |
| KYSE-150 | 3.67 | -2.70 |
| MCF-7 | 6.59 | -3.23 |
| 143-B | 7.34 | -4.75 |
| UACC-812 | 6.00 | -4.49 |
| SJSA-1 | 5.26 | -5.22 |
| PANC-1 | 12.00 | -3.16 |
| JeKO-1 | 6.35 | -9.80 |
| ZR-75-1 | 6.35 | -4.41 |

| | | |
|---|---|---|
| Capan-1 | 12.29 | -3.28 |
| CoC1-DDP* | 4.68 | -12.78 |
| SCC-4 | 7.43 | -3.09 |
| OE19 | 5.21 | -3.16 |
| A673* | 10.37 | -16.38 |
| NCI-1048* | 3.59 | -12.04 |
| AMO-1 | 6.12 | -5.82 |
| HT-29 | 6.51 | -4.24 |
| ES-2 | 4.96 | -4.41 |
| NCI-H209* | 10.59 | -15.84 |
| NCI-H1650 | 6.87 | -3.42 |
| HCC-4006 | 8.38 | -3.14 |
| HCT-15 | 9.00 | -4.96 |
| HCT-116 | 5.03 | -4.17 |
| NCI-H441 | 7.96 | -3.53 |
| KARPAS-299 | 4.74 | -4.94 |
| COLO-205 | 7.97 | -3.64 |
| KYSE 410 | 5.30 | -2.91 |
| BXPC-3 | 7.19 | -3.53 |
| Caki-1 | 6.06 | -5.80 |
| MS-751 | 6.22 | -1.73 |
| NCI-H929 | 4.87 | -5.36 |

Table 2. Replication dynamics images.

| Treatment | Representative images |
| --- | --- |
| DMSO | 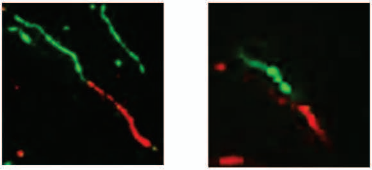 |
| HU | 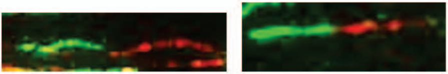 |
| ATRi | 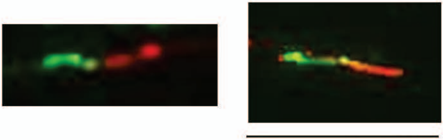 |
| HU+ATRi | 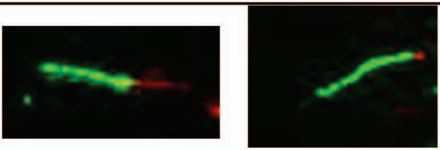 |
| RPAi | 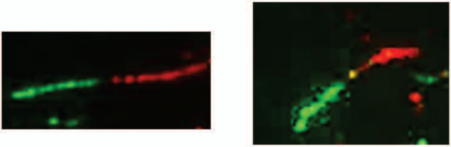 |
| HU+RPAi | 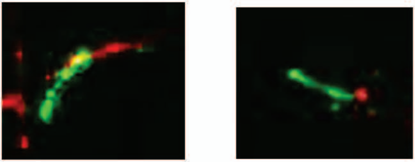 |
